# Knockdown of butyrylcholinesterase but not inhibition by chlorpyrifos alters early differentiation mechanisms in human neural stem cells

**DOI:** 10.1101/354308

**Authors:** Angela K. Tiethof, Jason R. Richardson, Ronald P. Hart

**Affiliations:** Joint Program in Toxicology, Environmental and Occupational Health Science Institute, Rutgers University, Piscataway NJ 08854 USA; Environmental Health Sciences, Robert Stempel School of Public Health and Social Work, Florida International University, Miami, FL 33199 USA; Department of Cell Biology & Neuroscience and the Human Genetics Institute of New Jersey, Piscataway, NJ 08854 USA

**Keywords:** Neural stem cell, butyrylcholinesterase, chlorpyrifos, Notch, HES5

## Abstract

Butyrylcholinesterase (BChE) is the evolutionary counterpart to acetylcholinesterase (AChE). Both are expressed early in nervous system development prior to cholinergic synapse formation. The organophosphate pesticide chlorpyrifos (CPF) primarily exerts toxicity through inhibition of AChE, which results in excess cholinergic stimulation at the synapse. We hypothesized that inhibition of AChE and BChE by CPF may impair early neurogenesis in neural stem cells (NSCs). To model neurodevelopment in vitro, we used human NSCs derived from induced pluripotent stem cells (iPSCs) with a focus on initial differentiation mechanisms. Over six days of NSC differentiation, BChE activity and mRNA expression significantly increased, while AChE activity and expression remained unchanged. CPF treatment (10 μM) caused 82% and 92% inhibition of AChE and BChE, respectively. CPF exposure had no effect on cell viability or the expression of differentiation markers HES5, DCX or MAP2. However, shRNA-knockdown of BChE expression resulted in decreased or delayed expression of transcription factors HES5 and HES3. BChE may have a role in the differentiation of NSCs independent of, or in addition to, its enzymatic activity.

## 1. Introduction

Chlorpyrifos (CPF) is a widely used organophosphorus class pesticide. Because of concerns relating to neurodevelopmental toxicity, residential and public pest management use has been eliminated [1]. Epidemiological studies suggest that CPF exposure is correlated with adverse neurodevelopmental effects involving cognition, behavior and fetal growth [2-4]. Animal studies have evaluated the developmental neurotoxicity (DNT) of CPF using endpoints including motor activity, cognition, emotion/anxiety and social interaction [5-10]. However, the mechanism(s) responsible for these effects have not been determined, particularly in human neuronal precursor cells, the likely target in developing brain.

In animal studies, daily administration of 5 mg/kg CPF to pregnant mice between gestation days (GD) 7.5-11.5 resulted in morphological changes, including thinning of CA1 and CA3 layers of the somatosensory cortex, enlargement of the dentate gyrus of the hippocampus and a decrease in the ratio of neurons and glia in the somatosensory cortex [11]. Similar morphological changes were found in juvenile rats after prenatal CPF exposure [12]. These finding were supported by recent epidemiological data which compared high and low prenatal exposure groups and found brain changes by MRI imaging, including frontal and parietal cortical thinning [13]. Therefore, exposure to CPF during sensitive periods of development may affect the development of the nervous system.

Inhibition of acetylcholinesterase (AChE) has long been considered the primary mechanism of CPF neurotoxicity [14]. However, AChE may have a role in neurogenesis through its enzymatic activity, by interactions with other cellular factors [15-18], or through an alternate activity as an aryl acylamidase [19].

Both AChE and the pseudocholinesterase, butyrylcholinesterase (BChE), are expressed early in embryonic development [20] and display characteristic spatial and temporal regulation [21]. In neural crest cells, BChE is expressed during mitosis, followed by increased expression of AChE during migration and differentiation [22]. BChE promotes proliferation prior to differentiation [23-25], while AChE may be involved in neurite outgrowth and cell adhesion [17]. Support for the role of AChE in neurite outgrowth is based on demonstration that blockage of the peripheral site of the enzyme changes neurite outgrowth density and may influence branching [26]. These non-classical roles for AChE and BChE have been reviewed previously [19,26,27].

Neural Progenitor Cells (NPCs), also known as neural stem cells (NSCs), are multipotent and can differentiate into neurons, astrocytes and oligodendrocytes [28]. The neural ectoderm, composed of NPCs, becomes two critical layers, the ventricular zone and the marginal zone, within the fluid-filled vesicle of developing forebrain. Cell proliferation occurs through vertical cleavage, which expands the NPC population in the ventricular zone and results in cellular migration and differentiation of one daughter cell. Therefore, in vitro differentiation of NSCs provides an opportunity to assess the potential neurotoxicity of environmental factors such as exposure to CPF on aspects of early neurogenesis, including proliferation and differentiation [29,30]. To model potential effects of CPF on early neural differentiation, we chose to use human NSC prepared from a pluripotent stem cell (iPSC) culture. Our goal was to characterize the NSC system, identify increased expression of both AChE and BChE, and then use mRNA markers of early commitment to neuronal differentiation to evaluate the effect of inhibition of cholinesterases by CPF. In addition, we examined whether BChE activity or gene expression was required during early neuronal differentiation.

## 2. Materials and Methods

### 2.1 iPSC and NSC Culture

Human foreskin fibroblasts were reprogrammed into induced pluripotent stem cells (iPSC) by retroviral overexpression of transcription factors [31] and were grown feeder-free [32,33]. iPSC were characterized and grown as described previously [33].

NSCs were prepared as described [34], except that Noggin was utilized to inhibit BMP pathways and promote differentiation to NSCs [35]. iPSCS were plated on BD Matrigel™ diluted in DMEM/F12 (BD Biosciences). Cells were fed every other day with medium consisting of 50% MTeSR (Stem Cell Technologies) and 50% Neural Basal Medium [NBM; Neurobasal^^®^^ Medium (Gibco), 2% B27^^®^^ Supplement, 1% Insulin-Transferrin-Selenium (Gibco), 1% N2 Supplement (Gibco), 2 mM L-Glutamine (Gibco), 0.5% Penicillin Streptomycin (Gibco) and 500 ng/mL Noggin (Peprotech)]. Five days after passage, medium was changed to 100% NBM with 500 ng/mL Noggin and fed every other day for 10 days. Cells were then passaged onto dishes coated with 20 µg/mL laminin (Sigma) in NBM without Noggin. At approximately 60-70% confluence, NSCs were switched to proliferation medium (see below) and grown until about 90% confluent.

NSCs were cultured in proliferation medium containing 20 ng/mL Recombinant Human FGFb (PeproTech) with 50% DMEM/F12 with Glutamax (Gibco), 50% Neurobasal^®^ Medium (Gibco) and 0.5X N2 Supplement (Gibco) and B27^®^ Supplement, minus vitamin A (Gibco). Cells were plated on 1:4 Matrigel™ diluted in DMEM/F12, and passaged with Accutase™ (Stemcell Technologies, Inc.) at approximately 40,000 cells/cm^2^ every 3-4 days.

### 2.2 NSC Differentiation

For differentiation, cells were plated in T75 flasks at 13,500 cells/cm^2^ and allowed to grow to confluence. On Day 0, cultures were switched to differentiation medium containing 10 ng/ml Brain Derived Neurotrophic Factor (BDNF, PeproTech) in Neurobasal plus N2 Supplement. Cells were harvested using Accutase^™^ at Day 0 (prior to initiation of differentiation), Day 2, Day 4 and 6. The cell pellet was resuspended in phosphate buffered saline (PBS) and split 1/3 for RNA and 2/3 for enzyme assays. Pelleted cells were rinsed well using PBS and residual PBS was aspirated prior to freezing at -70°C.

### 2.3 Chlorpyrifos Exposure

Cells were exposed to CPF 10 µM (Chem Service Inc.) starting on Day 0 of differentiation. Culture medium was changed every other day during scheduled collection time points. Control cultures contained 1 µL/mL of 100% ethanol.

### 2.4 Lentivirus Production and shRNA Knockdown

For BChE knockdown, the Mission^^®^^ shRNA TRC2 lentiviral vector plasmid was purchased from Sigma (TRC number TRCN0000427955, Clone ID NM_000055.2-439s21c1) along with non-target shRNA control plasmid SHC216. Addgene third generation packaging plasmids were transfected at a ratio of 4:2:1:1 of target vector, MDLg/RRE, VSVg-MD2g and RSV-REV, respectively into HEK293T cells. The culture medium was changed daily, harvested for virus on day 2 and 3 (at approximately 48 and 72 post transfection) and stored at 4 °C. The medium was then centrifuged at 200 X g to remove cellular debris, and virus concentrated by centrifuging at 25,000 rpm for 2 hours. The supernatant was removed and virus was resuspended in 180 µL DMEM/F12 overnight and stored at -70°C prior to use.

To quantify virus the Sigma-Aldrich Lentiviral Titer p24 ELISA assay protocol was used and the viral supernatants were assayed using the Retrotek HIV-1 p24 Antigen ELISA (0801111, Zeptometrix Corp.).

For knockdown experiments, cultures were seeded at a density of approximately 13,500 cells/cm^2^ in 6 well plate culture dishes and then infected the next day with 6 TU per cell in medium containing 8 µg/mL protamine sulfate (Sigma Aldrich). Medium was changed the next day to remove virus and protamine sulfate, and lysates for RNA were collected on Day 0 and Day 6 of differentiation.

### 2.5 Quantitative Real-time Polymerase Chain Reaction

RNA was isolated using the RNeasy^^®^^ Mini Kit (Qiagen) using on-column DNase treatment. 0.8 µg RNA was reverse transcribed using SuperScript ^™^ (Invitrogen) using random primers. Quantitative-PCR (qPCR) was performed using Power SYBR^^®^^ Green PCR Master Mix (Applied Biosystems). Relative mRNA expression was calculated using the 2^$^ ΔΔCt method [36] using TBP (TATA binding protein) as a normalizing gene. Primer sequences are listed in Table 1.

**Table 1:**
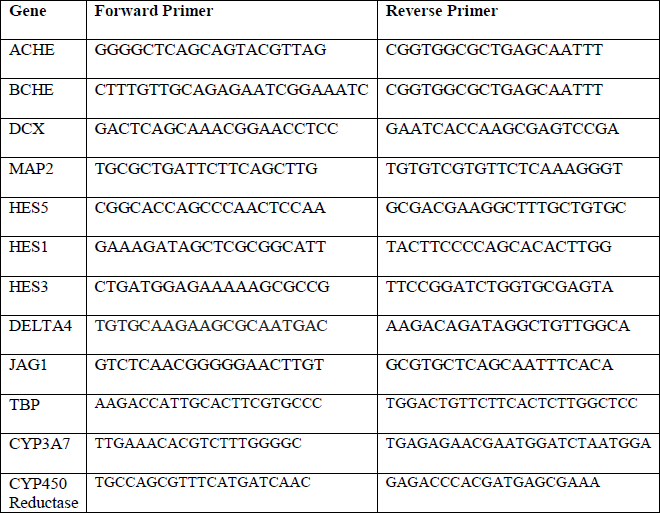
List of primers for qPCR. All sequences are 5’ to 3’.

### 2.6 Butyrylcholinesterase and Acetylcholinesterase Enzyme Assays

A 96 well microplate assay was developed from modification of the methods described previously [5]. Frozen cell pellets were re-suspended in 200-300 µL of 50 mM Tris buffer (pH 7.4 at 37 °C). Cell pellets were sonicated using a Labsonic ^^®^^ M (Sartorius Stedim Biotech). Between 15-30 µg of protein was assayed in duplicate spectrophotometrically, using 1 mM Acetylthiocholine iodide (Sigma-Aldrich) as the substrate for AChE activity and 2 mM SButyrylthiocholine iodide (Sigma-Aldrich) as the substrate for BChE activity. The chromagen 5,5’-dithio-bis(nitrobezoic acid) (DTNB, Sigma-Aldrich) was used at a final assay concentration of 0.3 mM, and 10 µM of eserine sulfate (Sigma-Aldrich) was used in duplicate parallel samples to correct for non-enzymatic substrate hydrolysis. The microplate containing sample, chromagen and inhibitor (if required) was pre-incubated for 10 minutes at 37°C with shaking. Substrate was added and the microplate was read at 412 nm for 20 min at 41 sec intervals with mixing. Specific activity was calculated as nmoles product formed per minute per mg protein using the extinction coefficient 14150 M^−1^cm^−1^.

### 2.7 Immunocytochemistry and Microscopy

NSCs were plated on poly-D-lysine (PDL, Sigma)/laminin (Sigma) coated 12 mm glass coverslips at 400,000 cells/well on Day 2 of differentiation and allowed to differentiate for 90 days. Cultures were fed with differentiation media containing 1 µg/mL laminin. On Day 90, cultures were then fixed for 30 minutes at room temperature in 4% paraformaldehyde, followed by an incubation for 1 hour with 4% normal goat serum and 0.01% triton x-100 in PBS to permeabilize and block. Cells were then incubated at 4°C overnight with the following antibodies: MAP2 (Microtubule associated protein 2, Millipore, 1:1000), Neuronal Class III β-Tubulin (TUJ1, Covance, 1:2000), Synaptophysin (Millipore, 1:500), vesicular glutamate transporter 1 (VGLUT1, Synaptic Systems, 1:500) and Glial Fibrillary Acidic Protein (GFAP, DakoCytomation, 1:1000), followed by removal of the primary antibody and triplicate PBS washes. The next day, cultures were incubated for 1 hour at room temperature with the following secondary antibodies: Alexa Fluor^^®^^ 488 conjugate (Goat anti-Rabbit IgG; Goat anti-Mouse IgG2A) and Alexa Fluor^®^ 594 conjugate (Goat anti-Mouse IgG1) at a 1:500 concentration. Nuclei were counterstained with DAPI (0.3 µg/ml, Roche) and rinsed well with PBS then distilled water prior to mounting.

### 2.8 Statistical Analysis

Experiments were performed at independent times and cell passages, and each independent differentiation was considered to be a biological replicate. To account for replicate differences, a one way ANOVA model was used to analyze changes in mRNA expression and enzyme activity. For CPF and lentivirus treated samples, paired t-tests were used for each time point to analyze expression differences between treatment and control. Data are presented as bar graphs with error bars representing mean ± standard error of mean (SEM).

## 3. Results

### 3.1 NSC Model of Early Neuronal Differentiation

NSCs were prepared from human iPSCs by switching from a proliferative medium containing FGF to a medium containing BDNF that favors differentiation (Fig 1A). During the initial six days of differentiation, cells transform from a homogenous appearance to one of clustered cell bodies, with outgrowth of processes visible between clusters (Fig. 1B). To assess NSC differentiation during this initial phase, three mRNAs were chosen as markers (Fig. 1C). HES5 was identified from an RNAseq analysis of similarly differentiating human ESC-derived NSCs [37]. HES5 is a repressive transcription factor which regulates neuronal differentiation, and, along with HES1, is an effector of the Notch pathway [38]. MAP2 is a cytoskeletal protein that associated with microtubules in neurons [39]. Doublecortin (DCX) is a microtubule-associated protein which is expressed by migrating neuroblasts during differentiation [40]. All mRNAs exhibited significant upregulation on Day 4 and 6, consistent with neuronal differentiation. Results indicate that NSC begin the process of differentiating into neurons within 2-6 days after BDNF addition and FGF withdrawal, and this provides a model for studying key mechanisms during initial commitment to neurons.

**Fig. 1:**
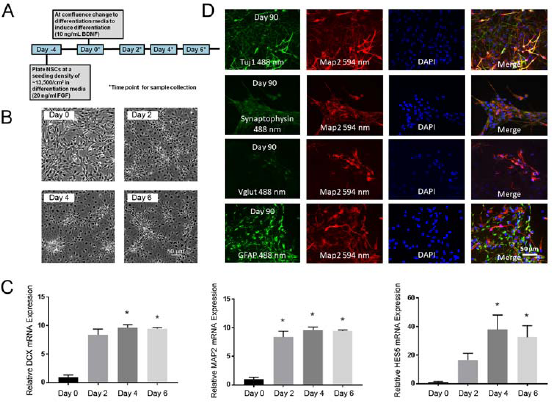
Characterization of Neural Stem Cell model of differentiation. (A) Overview of the experimental timeline of early differentiation. (B) Phase-contrast microscopy showing morphological changes of NSC in the first few days after induction of differentiation. (C) To assess early differentiation, the relative mRNA expression of the differentiation markers DCX, MAP2 and HES5 were determined by qPCR. All three markers increased significantly relative to Day 0. Results depict mean ± SEM (n=3-5) analyzed using a one-way ANOVA model to account for batch variation followed by Tukey multiple comparisons of means (*p < 0.05). (D) To confirm multipotency of NSC and that differentiation produces mature neurons and glia, cultures were fixed and stained immunocytochemically after 90 days of differentiation. All cultures consisted of cells positive for Tuj1, Synaptophysin, and VGlut (punctate staining, counterstained for MAP2) or GFAP (indicative of astrocytes, non-overlapping with MAP2).

To confirm the appropriate cellular identity of multipotent NSC, that they produce neurons and glia, we examined cultures differentiated for much longer periods of time for the production of mature neuronal morphologies and markers. In our previous experience with stem cell-derived NSC [41], ˜90 days normally produces robust expression of markers to evaluate neurons and astrocytes. Oligodendrocyte markers, such as MBP, were not evaluated. Cultures immunocytochemically stained after 90 days of differentiation indicated cells expressing the neuronal markers β_III_-tubulin (TuJ1), synaptophysin, and vesicular glutamate transporter (Vglut), each overlapping microtubule-associated protein 2 (MAP2), and the astrocytic marker GFAP, which did not overlap MAP2 (Fig 1D). Detection of diffuse synaptophysin immunoreactivity in the cytoplasm is consistent with neuronal expression prior to synaptogenesis, which would produce a more punctate staining. The large number of Vglut/MAP2 double-positive cells was consistent with the presence of glutamatergic neurons in these cultures. Some cells also stain positive for GFAP, which did not co-localize with MAP2, indicating astrocytes. GFAP- and MAP2-stained cells occurred in similar proportions, indicating a mixture of astrocytes and excitatory neuronal lineages. Expression of neuron and astrocyte markers at later time points indicate that NSC are multipotent and that the differentiation protocol is effective.

### 3.2 Butyrylcholinesterase mRNA Expression and Activity Progressively Increases During Differentiation

To determine whether AChE and BChE are regulated during early NSC differentiation, mRNA expression and enzyme activities were detected using qPCR and spectrophotometric assays, respectively (Fig. 2). During NSC differentiation, BChE mRNA expression and activity significantly increased during the initial 6 days. AChE mRNA and activity was unchanged relative to Day 0. This indicates that BChE mRNA expression and activity is upregulated along with other differentiation markers (HES5, DCX and MAP2), suggesting that BChE may play a role in early differentiation of NSCs.

**Fig. 2.**
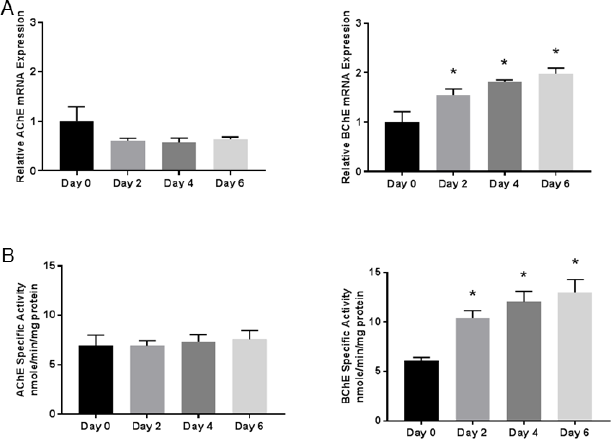
AChE and BChE activity and expression during early NSC differentiation. (A) The relative mRNA expression of the enzymes AChE and BChE were determined by quantitative real-time PCR. BChE mRNA expression increased significantly relative to Day 0 on all subsequent time points shown, while AChE was detected but unchanged over time. (B) AChE and BChE specific activity was assayed using a kinetic spectrophotometric assay. BChE specific activity increased significantly on all subsequent time points relative to Day 0, mirroring the mRNA expression. Data represent mean ± SEM (n=5-6), Data were analyzed using a one-way ANOVA model to account for batch variation followed by Tukey multiple comparisons of means (*p < 0.05). fic

### 3.3 Cholinesterase Inhibition by Chlorpyrifos Does Not Affect Neuronal Marker Expression

To test the hypothesis that CPF inhibition of cholinesterase affects early neurogenesis, NSC cultures were treated with 10 µM CPF. There was no effect on cell viability (as assayed with Alamar Blue^®^, not shown). CPF inhibited AChE and BChE activity 82% and 92%, respectively (Fig. 3A). This inhibition of activity did not change expression of AChE or BChE mRNAs relative to the control group (Fig. 3B). However, mRNA markers of neuronal differentiation (HES5, MAP2 and DCX) were not altered by CPF treatment (Fig. 3C).

**Fig. 3.**
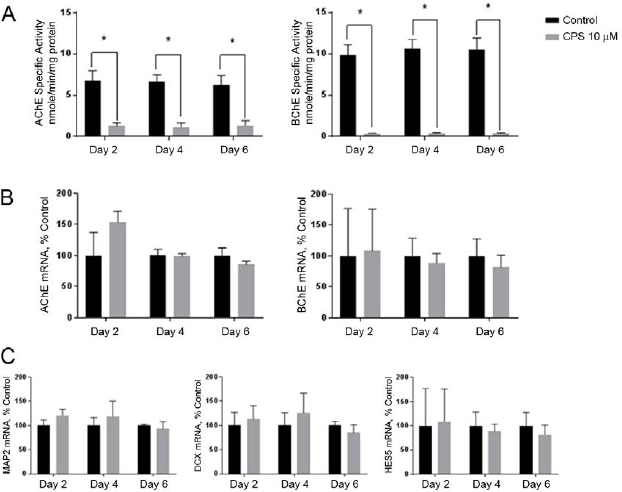
Inhibition of AChE and BChE by chlorpyrifos (CPF). Cells cultures were exposed continuously to 10 µM CPF starting on Day 0 (see Fig. 1a top right). (A) AChE and BChE specific activity was assayed using a kinetic spectrophotometric assay. Both AChE and BChE were inhibited, by 82% and 92% respectively. (B) The relative mRNA expression levels of the enzymes AChE and BChE and (C) the differentiation markers MAP2, DCX and HES5 were determined by quantitative real-time PCR. There were no significant differences between the control and 10 µM CPF exposure groups. Data represent mean ± SEM (n=3). Data were analyzed using paired t-tests for each time point to account for batch variation. (*p < 0.05).

To assess the potential for metabolic activation of CPF in NSC, CYP450 and CP450 reductase mRNAs were measured by qPCR. Both CYP3A7 and CYP450 reductase mRNAs were detected at all time points, with cycle threshold (C_t_) values of approximately 31.5 and 24.5, respectively, indicating a robust signal (results not shown). The likely presence of metabolizing enzymes and the observed cholinesterase activity inhibition indicates this model of NSC differentiation is likely to be capable of bioactivating CPF.

### 3.4 Knockdown of BChE mRNA Alters HES5 mRNA Expression

Since CPF inhibition of BChE activity did not affect markers of early differentiation, we considered whether BChE expression is required for normal differentiation. To test this alternative hypothesis that non-enzymatic functions of BChE regulate neurogenesis, NSCs were infected with lentivirus expressing BChE or negative control shRNA 72 hours prior to induction of differentiation on Day 0 (Fig. 4A). Knockdown of mRNA was calculated to be 72% (Fig. 4B). BChE knockdown resulted in a significant upregulation of AChE mRNA relative to control on Day 6 (Fig. 4C), no change in MAP2 (Fig. 4D) or DCX (Fig. 4E) mRNAs, but a down regulation of HES5 on Day 0 (Fig 4F). We conclude that BChE mRNA knockdown resulted in perturbed or delayed expression of the differentiation marker HES5.

**Fig. 4.**
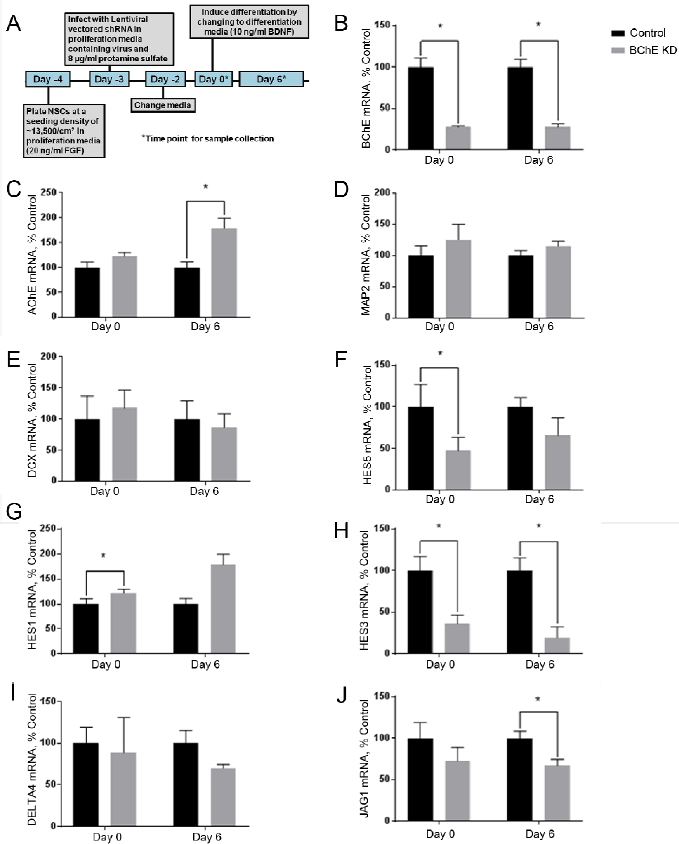
BChE shRNA knockdown. (A) Overview of the experimental timeline of differentiation. The relative mRNA expression levels of the enzymes (B) BChE and (C) AChE and the differentiation markers (D) MAP2, (E) DCX, (F) HES5, (G) HES1, (H) HES3, the Notch ligands n. (I) DELTA4 and (J) JAG1 were determined by qPCR. BChE knockdown was calculated to be 72%. AChE expression was significantly increased on Day 6 relative to control. Of the early differentiation markers, only HES5 was significantly decreased on Day 0 of differentiation, but components of the NOTCH signaling pathway (HES1, HES3, and JAG1) were affected by BChE knockdown. Data represent mean ± SEM (n=4). Data were analyzed using paired t-tests for each time point to account for batch variation. (*p < 0.05).

Because HES5 is regulated by the Notch pathway, we assayed the mRNA expression of additional Notch pathway genes HES1, HES3 and the Notch ligands JAG1 and DELTA4 following BChE knockdown (Fig 4G-J). HES1 was upregulated on Day 0 relative to the control (Fig. 4G). The Day 0 sample was taken prior to BDNF addition and FGF withdrawal. HES3 was downregulated both on Day 0 and Day 6 (Fig. 4H) and the Notch ligand DELTA4 was unaffected (Fig. 4I) but JAG1 was down regulated on Day 6 (Fig. 4J). Results indicate that BChE mRNA knockdown, but not inhibition by CPF, produced changes in the mRNA expression of several developmental and Notch pathway genes, suggesting that BChE likely has a non-enzymatic role in NSC differentiation.

## 4. Discussion

We established an iPSC-derived NSC human cell culture model of early neuronal differentiation as a model for testing neurotoxicity in a human neural progenitor. This system was used to probe the DNT of CPF and to assess the function of BChE early in neurogenesis. The NSC model consists of both a proliferative phase and a differentiation phase, with differentiation characterized by morphological changes and increased expression of the neuronal structural markers MAP2, DCX and the transcription factor HES5. These three markers were chosen from a comprehensive RNAseq analysis of NSC differentiation in an earlier study [41] as preliminary indicators for changes occurring during the initial phase of neurogenesis. Differentiation is induced by the withdrawal of pro-mitotic FGF and the addition of differentiation-promoting BDNF. Much later in the process, after 90 days, cells stain positive for both neuronal markers, including TuJ1, synaptophysin, Vglut, and MAP2, as well as the non-overlapping astrocytic marker GFAP (Fig 1C), as direct proof that the NSC cultures were multipotent. This model also displayed expression of both CPF targets BChE and AChE, although only BChE mRNA and activity was increased during the six days of differentiation. BChE is known to be expressed at higher levels prior than AChE in early embryonic development, and, in neural crest cells, BChE is expressed during mitosis and AChE is expressed during migration and through differentiation [21]. This characteristic early expression of BChE during early differentiation is consistent with the hypothesis that BChE may have a role in early NSC differentiation.

To determine whether the catalytic activity of AChE and BChE is essential for early NSC differentiation, cells were treated continuously with a concentration of CPF sufficient to inhibit AChE and BChE activities, but not enough to cause overt cytotoxicity. Interestingly, we were able to detect expression of both the CYP3A7 and CYP450 reductase mRNAs in NSC model, likely indicating the capacity to convert CPF to its more toxic oxon.

Others have devised strategies to test neurotoxicity in cellular models of neural progenitors or neurons. In a study using NSCs derived from human umbilical cord blood to model DNT, minimal toxicity was observed at 10 µM CPF, although cells differentiated more towards an astrocytic phenotype and showed partial decrease in viability [42]. Another recent study used a commercial, immortalized human cell line to test epigenetic effects of CPF in both proliferating and differentiating NSCs, and toxic effects were not detected at concentrations below 57 µM [43]. Neither study quantified cholinesterase activity. Another study used adipose-derived stem cells to model CPF toxicity [44], but used a much higher dose of CPF (500 μM), finding decreased viability in differentiating cells. One study examined differentiation outcomes in embryonic stem cell-derived NPCs, finding reduced proportions of glial cells upon exposure to 10-30 μM CPF [45]. Others used neuron-like cell lines including N2a and PC12 cells, identifying cellular effects such as neurite retraction [46,47], which would reflect events much later than the initial NSC differentiation modeled here. Our goal was to identify specific mechanisms affected early in NSC differentiation, since these events are likely to cause changes in differentiation choice, such as the reduced glial differentiation [44], or changes in neuronal circuitry due to shifted differentiation patterns.

CPF inhibition of both AChE and BChE activities, however, did not alter expression of the early differentiation markers MAP2, DCX, or HES5 (Fig. 3C). The dose of 10 μM CPF was chosen due to the lack of observed cytotoxicity and to mimic lower levels of exposure likely to be found in exposed subjects during fetal development. However, it is within the range of doses found to alter eventual balance of mature cell types produced later in development [45]. We conclude that CPF did not affect initial molecular mechanisms during early neurogenesis from progenitor cells.

One possibility was that neither AChE nor BChE are required for early neurogenesis. Since we observed an increase in mRNA and activity for BChE during this phase (Fig. 2), we considered that BChE protein may be required for differentiation of NSCs. To test this, we used lentiviral-encoded shRNA to knockdown BChE. BChE knockdown reduced expression of HES5 mRNA by over 50%, but did not affect MAP2 or DCX mRNAs, suggesting a selective effect. Because HES5 is known to be regulated by the NOTCH signaling pathway we also determined that HES1 and JAG1 were altered (Fig. 4G,J). Interestingly, prior to the initiation of differentiation, BChE knockdown reduced HES3 (Fig. 4H) and HES5 (Fig. 4F) mRNA levels, and increased HES1 (Fig. 4G) mRNA, suggesting that endogenous BChE expression affects HES signaling in NSC. The repressor-type basic helix-loop-helix (bHLH) genes HES1, HES3 and HES5 play a critical role in NSC differentiation and maintenance [48]. Activating bHLH target genes, including Mash1, Math and Neurogenin, promote neurogenesis, whereas the repressor bHLH genes promote gliogenesis, which are formed later in neural stem cell differentiation.

HES1 and HES5 are regulated by the Notch pathway [49] but HES3 is not [50]. HES1 and HES5 can compensate for each other, although HES1/HES5 double knockout mice show decreased maintenance of glial cells and increased early neurogenesis [48]. Triple knockout HES1/HES3/HES5 mice have a more severe phenotype, which results in premature differentiation of stem cells in neurons, and a loss of other cell types including astrocytes and oligodendrocytes [51]. Because HES5 expression was decreased with BChE knockdown, mediation by Notch pathway proteins is likely. The notch ligand JAG1 was decreased on Day 6 of differentiation. JAG1 is involved postnatal and adult neurogenesis in the dentate gyrus [52], and astrocytes have been shown to decrease neurogenesis though Notch/JAG1 signaling [53].

Based on these results, we propose that BChE may interact with other proteins in NSC and early during initial NSC differentiation through a non-enzymatic mechanism. It has been suggested that ChE may act as cell adhesion molecules (CAMs). CAMs that function in nervous system development have an extracellular domain which displays sequence homology to ChEs [54], and the X-ray crystallography structure of human BChE had structural homology with neuroligins [55]. Knockdowns indicated that BChE antisense treatment reduced proliferation and increased differentiation and apoptosis in chick retinae [23,25] and in a rat oligodendroglia cell line [24]. BChE antisense transfection reduced proliferation, increased protein kinase C (PKC) and pushed the cells towards an astrocytic phenotype. Finally, knockdown of BChE in R28 cells, a retinal rat cell line with pluripotent characteristics increased AChE expression, and this “counter-regulation” was tightly controlled through transcription factors c-fos and P90RSK1 [56]. BChE knockdown perturbed PKC and ERK signaling suggesting that coordinated ChE expression is involved in cell fate determination. Further research is required to fully elucidate the relative roles of AChE and BChE protein in NSC differentiation and whether this process may be targeted by environmental toxicants such as CPF to disrupt NSC differentiation.

In conclusion, we established a human iPSC-derived NSC model of early differentiation to probe the DNT of CPF. Although CPF inhibited the enzymatic activity of both AChE and BChE, no effects on NSC toxicity or on the mRNA expression of early differentiation markers HSE5, MAP2 and DCX were found. On the other hand, shRNA knockdown of BChE resulted in decreased mRNA expression of HES5, and changes in the expression of the related genes HES3, HES1 and JAG1. These results suggest a novel and potentially non-enzymatic role of BChE in NSC differentiation mechanisms through regulation of HES genes, possibly through the Notch pathway.

## Acknowledgements

Supported by a pilot award from P30 ES005022 and by R01 ES026057 (JRR and RPH).

